# Increased production of the extracellular polysaccharide Psl can give a growth advantage to *Pseudomonas aeruginosa* in low-iron conditions

**DOI:** 10.1101/355339

**Authors:** Jaime Hutchison, Karishma Kaushik, Christopher A. Rodesney, Thomas Lilieholm, Layla Bakhtiari, Vernita D. Gordon

## Abstract

In infections, biofilm formation is associated with a number of fitness advantages, such as resistance to antibiotics and to clearance by the immune system. Biofilm formation has also been linked to fitness advantages in environments other than *in vivo* infections; primarily, biofilms are thought to help constituent organisms evade predation and to promote intercellular signaling. The opportunistic human pathogen *Pseudomonas aeruginosa* forms biofilm infections in lungs, wounds, and on implants and medical devices. However, the tendency toward biofilm formation originated in this bacterium’s native environment, primarily plants and soil. Such environments are polymicrobial and often resource-limited. Other researchers have recently shown that the *P. aeruginosa* extracellular polysaccharide Psl can bind iron. For the lab strain PA01, Psl is also the dominant adhesive and cohesive “glue” holding together multicellular aggregates and biofilms. Here, we perform quantitative time-lapse confocal microscopy and image analysis of early biofilm growth by PA01. We find that aggregates of *P. aeruginosa* have a growth advantage over single cells of *P. aeruginosa* in the presence of *Staphylococcus aureus* in low-iron environments. Our results suggest the growth advantage of aggregates is linked to their high Psl content and to the production of an active factor by *S. aureus*. We posit that the ability of Psl to promote iron acquisition may have been linked to the evolutionary development of the strong biofilm-forming tendencies of *P. aeruginosa*.

## Introduction

Biofilms are sessile communities of interacting microbes that are embedded in a matrix dominated by extracellular polysaccharides (EPS) that holds bacteria in place [1–4]. For infections, being in a biofilm confers advantages on the constituent bacteria by helping them resist antibiotics and evade the immune system [5–8]. *Pseudomonas aeruginosa*, is an opportunistic pathogen that forms biofilms in many scenarios: in the lungs of patients with Cystic Fibrosis and Chronic Obstructive Pulmonary Disease; in chronic wounds, especially but not exclusively in patients with diabetes; and on implanted medical devices [7, 9–12]. However, *P. aeruginosa* has its primary home, and much of its evolutionary history, on soil and plants [13–15]. Other researchers have shown that, in such natural environments, aggregation might help protect against predation and chemical attack, and allow beneficial inter-cellular cooperative behaviors [16–20]. However, little is known about how EPS-driven aggregation may be linked to resource acquisition and resulting growth advantage and to the chemical properties of specific EPS types. This gap in knowledge prevents the development of ecology-based strategies for thwarting harmful biofilms or promoting beneficial biofilms.

In this work, we find that when iron is a limiting resource, multicellular aggregates of *P. aeruginosa* have a growth advantage over *P. aeruginosa* single cells that is linked to iron acquisition and to the presence of *Staphylococcus aureus. S. aureus* is a Gram-negative bacterium that is often found as a co-pathogen in *P. aeruginosa* biofilm infections, and can also be thought of as a token representation of the diverse microbial populations in the ecologies in which *P. aeruginosa* evolved. We also find that the enhanced growth of aggregates is linked to aggregates having higher amounts of the extracellular polysaccharide Psl than have single cells. Therefore, we infer that the need to acquire and use iron, as a vital growth substrate, could result in a competitive advantage to the production of Psl, which both binds iron [21] and also promotes intercellular aggregation and biofilm formation.

In contrast to the present work, we have previously shown that, in monoculture biofilms of *Pseudomonas aeruginosa* grown in flow cells, aggregates can have a growth advantage over single cells *when the competition for growth resources is high* [22]. We attributed this effect to cells at the top of the aggregate having greater access to growth resources, which are supplied diffusively from far above the floor of the flow cell, where the single cells are attached [22]. Thus, the growth advantage we found for aggregates in our previous work is a direct consequence of both the three-dimensional structure of aggregates and the spatial structure of the environment. Our earlier work also showed that when competition for resources is low, aggregates have a growth disadvantage compared with single cells [22]. This arises because cells in the interior of the aggregate have lower access to growth resources than do single cells. Although this disadvantage is present regardless of the level of competition for resources, it puts the aggregates at an *overall* disadvantage compared with single cells only when the competition for resources is low.

Unlike our earlier work comparing aggregates with single cells in monoculture [22], the iron- and Psl-linked advantage for aggregates that we investigate here does not depend on competition or on vertical structure of the aggregates or the environment.

## Materials and Methods

### Bacterial strains

All strains of *Pseudomonas aeruginosa* were in the background of the PA01 lab strain and were obtained from Colin Manoil at the University of Washington [23]. PA01 biofilms and aggregates are Psl-dependent and -dominated [24]. To allow visualization by confocal fluorescence microscopy, wild-type (WT) PAO1 strains were tagged with mCherry *via* Tn7 transformation and Δ*pel* PAO1 strains were tagged with green fluorescent protein (GFP) on plasmid pMRP9-1. As a prototypic Gram-positive organism, *Staphylococcus aureus* strain MN8 expressing yellow fluorescent protein (YFP) *sar* P_1_-*yfp*_*10B*_ on plasmid pJY209 was used for all experiments containing *S. aureus*, in monoculture or in co-culture with *P. aeruginosa* (this plasmid was a gift from Marvin Whiteley, University of Texas, Austin).

## Bacterial growth media and culture conditions

### Growth before microscopy

All strains were streaked onto Luria-Bertani (LB) agar plates (Fisher Scientific, USA) and incubated overnight at 37°C *P. aeruginosa* and *S. aureus* colonies were inoculated into separate tubes of 4 mL LB broth (Fisher Scientific, USA), and grown overnight with shaking at 37°C. The optical density (OD) of the overnight cultures was measured with a spectrophotometer (Genesys, USA). Overnight shaken cultures of *P. aeruginosa* naturally contain both multicellular aggregates and single cells [25], and aggregates analyzed in the experiments formed spontaneously during overnight growth.

### Growth under the microscope

For microscopy, the medium used was M9 minimal media (Serva, Germany) with 10% v/v A10 phosphate buffer and a final concentration of 0.3 mM filter-sterilized glucose (Fisher Scientific, USA). This medium will be referred to as “M9+glucose” in the rest of this paper. Immediately prior to the start of the flow cell experiment, *P. aeruginosa* and *S. aureus* cultures were mixed in desired ratios (1:1 if not explicitly stated otherwise) into 10 mL of M9+glucose to achieve the desired combined final OD (0.1 for high-density, 0.01 for medium-density and 0.001 for low-density).

For experiments with explicitly-added iron, ferrous sulfate, FeSO_4_ (Sigma Aldrich, USA), was dissolved in distilled water, filter sterilized, and added to M9+glucose for a final concentration of either 0.05, 5, or 100 μ-molar.

For experiments with an iron chelator, a stock solution of de-ferrated ethylene diamine diorthohydroxyphenyl acetic acid (EDDHA) at a concentration of 50 mg/mL in 1N NaOH, adjusted to a pH of 9, was prepared and stored at 4°C in plastic tubes (deferrated EDDHA was a gift from Shelley Payne, University of Texas, Austin). EDDHA primarily chelates the ferric form of iron, though it can also chelate small amounts of ferrous iron. EDDHA was added to M9+glucose at a final concentration of 32 μMolar and 320 μMolar, representing low and high concentration of iron chelator.

### Flow cell experiments

We used a standard flow cell system as previously described with certain modifications [22, 26, 27]. A three-chamber flow cell was filled with pre-warmed 37°C M9+glucose. Pre-warming the media reduces the likelihood of bubbles entering the sample chamber. Individual flow chambers were inoculated by directly injecting 200-300 μL of bacterial suspension using a 24 gauge needle and 3 mL syringe. The small puncture in the silicone tubing caused by the injection process was sealed with silicone sealant. After inoculation, the flow cell was left static (without flow) for 1 hour to allow bacteria to attach to the coverslip. After that, laminar flow at 3 mL/hour was provided by a peristaltic pump (Watson-Marlow) for the entire duration of the experiment.

## Imaging

Biofilms were imaged using an Olympus IX81 inverted confocal microscope (Olympus, Japan) with a 60x oil-immersion objective. The stage was enclosed by an incubation chamber heated to 37 °C, which is human physiological temperature. Image capture was controlled by FV10-ASW version 3.1 software. Areas of interest were selected by identifying suitable *P. aeruginosa* aggregates paired with regions of *P. aeruginosa* single cells that were each ~500 μm away from the corresponding aggregate (Fig 1). Single-cell regions were displaced from their corresponding aggregate in a direction perpendicular to the direction of media flow, so that the aggregate and singlecell region were at the same distance from the media inlet, and therefor were in areas with approximately the same concentration of growth resources such as carbon source, oxygen, and iron, and of metabolic byproducts. A displacement of ~500 μm subtends approximately 10% of the width of the ~5mm flow channels and is ~10× greater than ~10 μm aggregate diameter. This was chosen to achieve spacing between observed regions such that the single-cell region was far from, and unperturbed by, the aggregate, but the perpendicular distance from either region to the wall of the flow cell was not materially changed, as regions of observations were chosen to be roughly centered on the width of the flow chamber.

**Fig 1.**
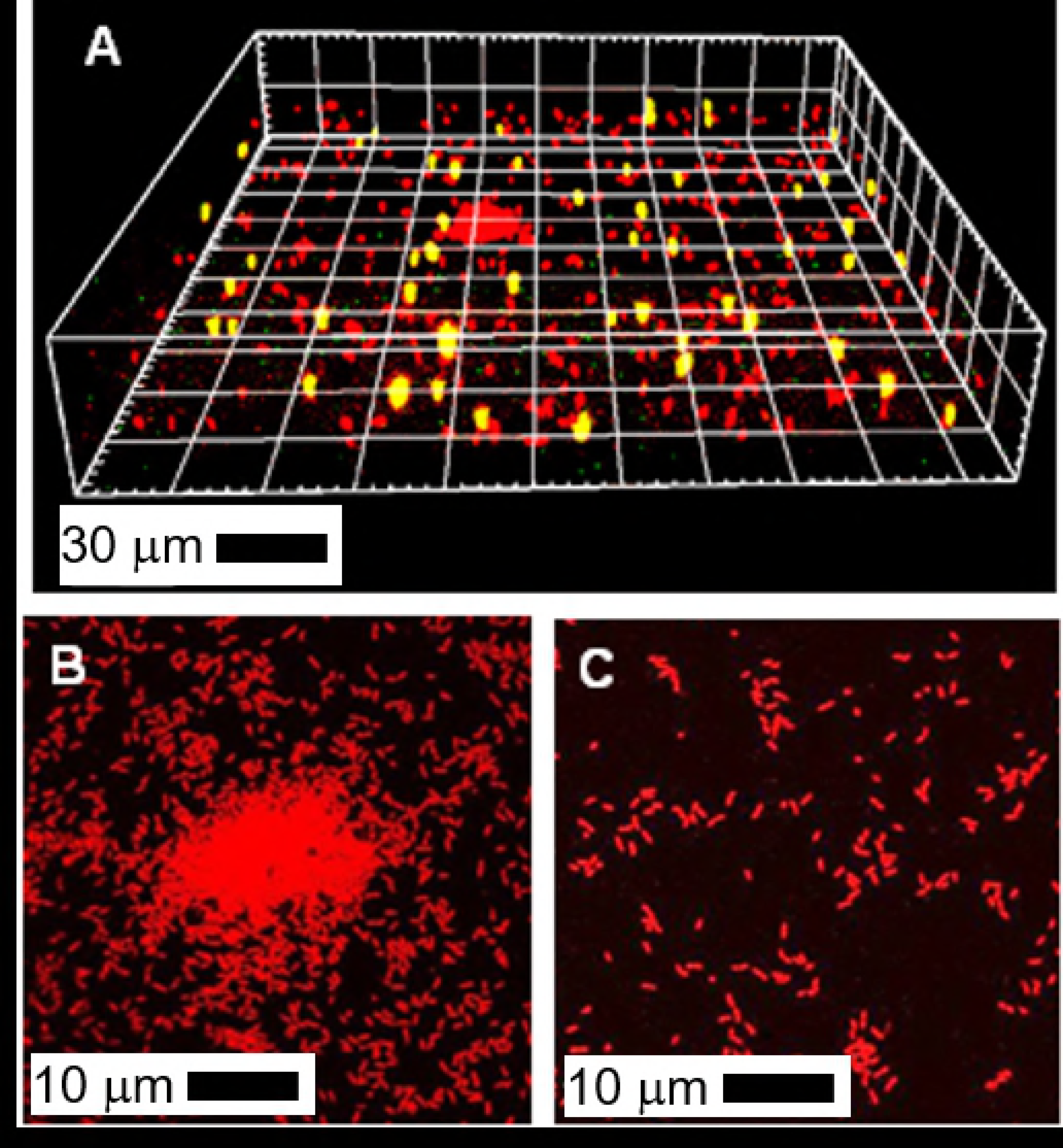
Confocal microscopy reveals the position, density, and (with timelapse microscopy) growth of *P. aeruginosa* single cells and multicellular aggregates and *S. aureus* bacteria. (A) A 3-D volume of a region seeded with a mixture of *S. aureus, P. aeruginosa* single cells, and a *P. aeruginosa* aggregate. *P. aeruginosa* (fluorescent with mCherry) are shown in red and *S. aureus* (fluorescent with YFP) appear yellow. (B) A 2-D projection of a multicellular *P. aeruginosa* aggregate. (C) Single cells of *P. aeruginosa* at 500 mm distance from the aggregate in panel (B). Single cells are always examined at a fixed distance of 500 mm from the aggregate to which they are compared, in a direction perpendicular to the direction in which medium is flowing along the length of the flow cell. Since the availability of growth substrates such as oxygen decrease with the direction of flow, comparing the growth of aggregates and single cells at the same position along the length of the flow cell normalizes for differences in growth rates arising from different concentrations of growth substrate.

An aggregate of *P. aeruginosa* was defined as a clump of at least 10 cells adhering to each other and extending vertically above the surface of the flow cell. A programmable motorized stage was used to cycle between multiple locations (typically n=5-20) in each chamber throughout the experiment. At each position of interest, Z-stacks were taken every 1.5 hours for the first 9 hours of biofilm growth.

## Data Analysis

Images were processed using ImageJ (National Institutes of Health, USA). For coculture experiments, the first step in processing was to account for bleed-through of the *S. aureus* fluorescence into the *P. aeruginosa* images by subtracting the yellow *S. aureus* channel from the green or red *P. aeruginosa* channel. Next, images containing aggregates were cropped manually to separate aggregates and their descendants from single cells and their descendants. *P. aeruginosa* biomass at time=0 (N_0_) and time=9 (N) hours was then determined by Matlab (MathWorks, USA) using software code that we wrote for earlier work [22, 27].

We then calculated N/N_0_ for the *P. aeruginosa* aggregate 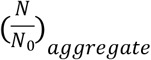 and N/N_0_ for the *P. aeruginosa* single cells 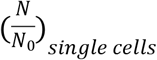 for each pair of images. Thus, one aggregate is associated with one area of single cells ~500 μm away, and these regions are paired for analysis. The relative fitness *f* was calculated as the ratio 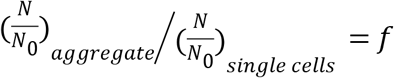.

Thus, a relative fitness greater than one indicates that aggregates are fitter than single cells, a relative fitness less than one indicates that single cells are fitter than aggregates, and a relative fitness of one indicates no fitness difference between aggregates and single cells.

## Statistics

We checked for statistically significant differences between two data sets using the two-tailed Student t-test. For comparison of a data set to unity, we used the one-sample t-test, where the value for the null hypothesis was 1. We take *p*-values ≤ 0.05 to indicate statistical significance, and *p*-values are indicated above each bar on plots as (*) *p* ≤ 0.05, (**) *p* ≤ 0.01, (***) *p* ≤ 0.001, or (ns) *p* > 0.05.

## Results

### Co-culture with *S. aureus* increases the fitness of *P. aeruginosa* aggregates over *P. aeruginosa* single cells, in a competition-independent way

In this subsection, we assess the effect of co-culture with *S. aureus* and the effects of net cell density on the growth of *P. aeruginosa* aggregates and of *P. aeruginosa* single cells. For the experiments discussed in this sub-section, the ratio of *P. aeruginosa* to *S. aureus* in the co-culture innocula, as measured by the OD of each culture before mixing, was held constant at a nominal 1:1. By varying the total cell density (*P. aeruginosa* + *S. aureus*) of the inoculum to correspond to ODs of 0.1 (high), 0.01 (medium) and 0.001 (low), we varied the seeding density of single cells and thus the level of competition for growth resources on the coverslip surface of the flow cell [22]. We also seeded with monoculture innocula consisting of *P. aeruginosa* alone at ODs of 0.1 (high), 0.01 (medium) and 0.001 (low), which largely corresponds to the competition measurements done in our previous work [22]; the primary difference is that, in the present experiments, the lateral position along the flow cell was kept constant as pairs of aggregates and singlecell regions were compared.

In monoculture, we find that aggregates of *P. aeruginosa* grow better than single cells of *P. aeruginosa* at high cell density; aggregates grow worse than single cells at low cell density; growth of aggregates and single cells is comparable at medium cell density. These finding agrees with our earlier publication showing that *P. aeruginosa* aggregates can have a competition-dependent growth advantage over *P. aeruginosa* single cells [22]. We also find that co-culture with *S. aureus* does not harm the growth of *P. aeruginosa* (Fig 2). Indeed, co-culture with *S. aureus* slightly benefits *P. aeruginosa* growth, although in most cases the increase in growth is not statistically significant (Fig 2).

**Fig 2.**
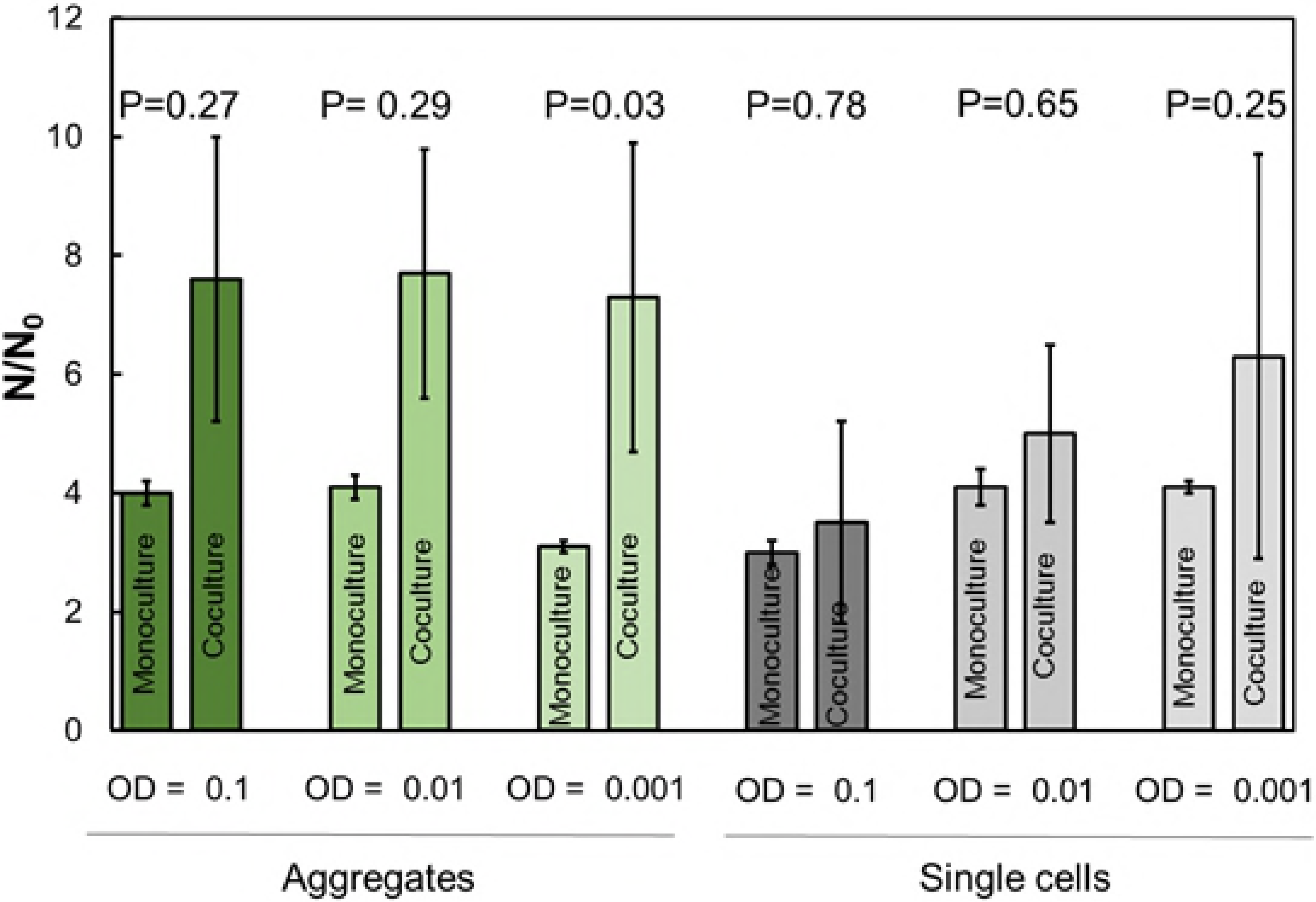
Co-culture with *S. aureus* does not harm the growth of *P. aeruginosa*. Growth is measured as biomass after nine hours of growth (N) divided by initial biomass (N_0_). Data are paired by monoculture *P. aeruginosa* and *P. aeruginosa* in co-culture with *S. aureus* for each net cell density (OD) examined. P-values compare each pair of bars. The growth of aggregates (in green) is measured separately from the growth of single cells. The growth of *P. aeruginosa* aggregates is greater when *S. aureus* is present in co-culture, but the difference in growth rate is only statistically significant in one case, at low OD (OD=0.001). The growth of *P. aeruginosa* single cells is marginally greater when *S. aureus* is present in co-culture, but these differences are not statistically significance. Error bars are standard error of the mean.

However, we find that co-culture with *S. aureus* enhances the growth of *P. aeruginosa* aggregates more than it enhances the growth of single cells. At high and medium cell density, in co-culture with *S. aureus*, the growth of *P. aeruginosa* aggregates is greater than the growth of *P. aeruginosa* single cells and these differences are statistically significant; at low cell density, in co-culture with *S. aureus*, the growth of *P. aeruginosa* aggregates is greater than the growth of *P. aeruginosa* single cells but this difference is not quite statistically significant (Fig 3). The fitness advantage of aggregates over single cells is not statistically different for the three cell densities studied, and thus does not depend on competition (Fig 3). This is strikingly different from our earlier work, in *P. aeruginosa* monoculture, in which aggregates had a fitness advantage only at high competition and had a fitness <u>disadvantage</u> at low competition [22].

**Fig 3.**
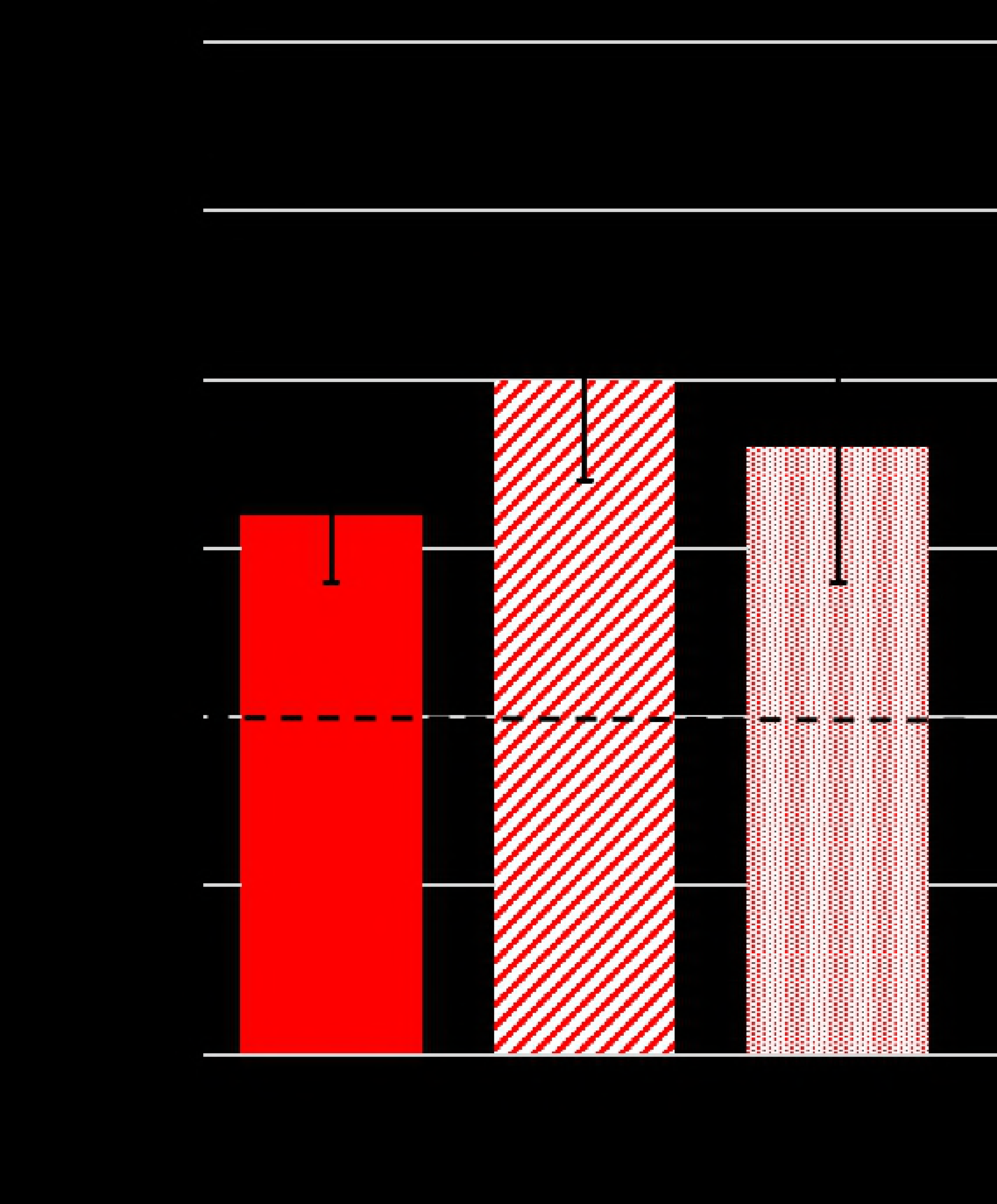
In co-culture with *S. aureus*, multicellular aggregates of *P. aeruginosa* are fitter than single cells of *P. aeruginosa*. We quantify the growth advantage of aggregates using the relative fitness measure defined in Materials and Methods, such that a relative fitness value *f* > 1 means that aggregates are fitter than single cells; *f* =1, indicating no fitness difference between aggregates and single cells, is indicated with a dashed line. The average value of the relative fitness is 1.6± 0.4 for high cell density (OD = 0.1), 2.0 ± 0.3 for medium cell density (OD = 0.01), and 1.8 ± 0.4 for low cell density (OD = 0.001). To determine statistical significance for each density, we used a null hypothesis that *f* =1 - i.e., that aggregates and single cells grow at equal rates (shown by the dashed line). High- and low-density cases have *p*-values of 0.01, indicating statistical significance. To assess the effect of cell density, i.e. competition, on the relative fitness of aggregates, we compare the relative fitness of aggregates across each pair of densities, with a null hypothesis that the *f* is the same in all cases; the resulting p-values, indicated by dotted lines comparing pairs of conditions, show no significant difference caused by varying density.

Our previous work has shown that under monoculture conditions, *P. aeruginosa* multicellular aggregates are more fit than corresponding single cells at high levels of competition for nutrients (where cell density is a proxy for competition) [22]. Thus, our results with coculture could be explained if *S. aureus* were simply a better competitor for resources than *P. aeruginosa*. To investigate this possibility, we grew *S. aureus* in monoculture, both as planktonic cells in shaking culture and in the biofilm mode in a flow cell. We monitored bacterial growth under both conditions. Under planktonic monoculture conditions, we find that *S. aureus* does not grow well in M9+glucose medium (Supplementary Fig 1A). This contrasts with monocultures of *P. aeruginosa* grown under the same conditions, which display typical growth dynamics with entry into the exponential phase approximately 20-30 hours after innoculation. In flow cells with M9+glucose medium, *S. aureus* cultures also display poor growth and very little biofilm even after 12 hours of growth (Supplementary Fig 1B). Thus, we conclude that it is unlikely that a poorly-growing culture of *S. aureus* would be a better competitor for resources than heartily-growing *P. aeruginosa*, and therefore that the dependence of aggregate relative fitness on local *S. aureus* does not arise from competition.

Thus, we find that co-culture with *S. aureus* increases the relative fitness of multicellular aggregates of *P. aeruginosa* and erases the dependence of relative fitness on competition that characterizes the monoculture system.

### The relative fitness of *P. aeruginosa* aggregates depends on the local ratio of *S. aureus* to *P. aeruginosa*

To explore the role that *S. aureus* plays in the relative fitness of multicellular aggregates of *P. aeruginosa*, we varied the initial sample-wide ratio of *S. aureus* to *P. aeruginosa* when combining them prior to inoculation (*P. aeruginosa* < *S. aureus*, *P. aeruginosa* ~ *S. aureus*, and *P. aeruginosa* > *S. aureus*) to a final OD of 0.01. As measured by OD of the samples pre-mixing, the *P. aeruginosa* < *S. aureus* samples had *S. aureus* concentration four times that of *P. aeruginosa*, the *P. aeruginosa* > *S. aureus* samples had *P. aeruginosa* concentration four times that of *S. aureus*, and the *P. aeruginosa* ~ *S. aureus* samples had *P. aeruginosa* concentration the same as that of *S. aureus*. We find that the relative fitness of aggregates does not depend on the sample-wide ratio of the two species.

To assess how the relative fitness of *P. aeruginosa* aggregates depends on the <u>local</u> concentration of *S. aureus*, for each field of view (roughly 250 μm × 250 μm), we measured the initial *S. aureus* biomass and the initial *P. aeruginosa* biomass to determine the initial local ratio of *S. aureus* to *P. aeruginosa*; only single cells in the fields of view were used for these measurements. We then binned the resulting relative fitness of *P. aeruginosa* aggregates according to the initial local ratio of *S. aureus* to *P. aeruginosa*, with bins as follows: *P. aeruginosa*: *S. aureus* < 1; 1 < *P. aeruginosa*: *S. aureus* < 10; *P. aeruginosa*: *S. aureus* > 10 (Fig 4). We find that when initial local concentrations of *S. aureus* > *P. aeruginosa*, the average relative fitness of aggregates is 2.5 ± 0.4. When the initial local concentrations are such that *S. aureus* ~ *P. aeruginosa*, the average relative fitness is 1.8 ± 0.3. These values for relative advantage are both statistically significant. When the initial concentrations of *S. aureus* < *P. aeruginosa* (*i.e*. when approaching a monoculture of *P. aeruginosa*) there is no statistically-significant relative fitness advantage for aggregates. Thus, we find that the relative fitness of *P. aeruginosa* multicellular aggregates depends on the initial **local** ratio of the two species, increasing as the local population contains a higher proportion of *S. aureus* than *P. aeruginosa*.

**Fig 4.**
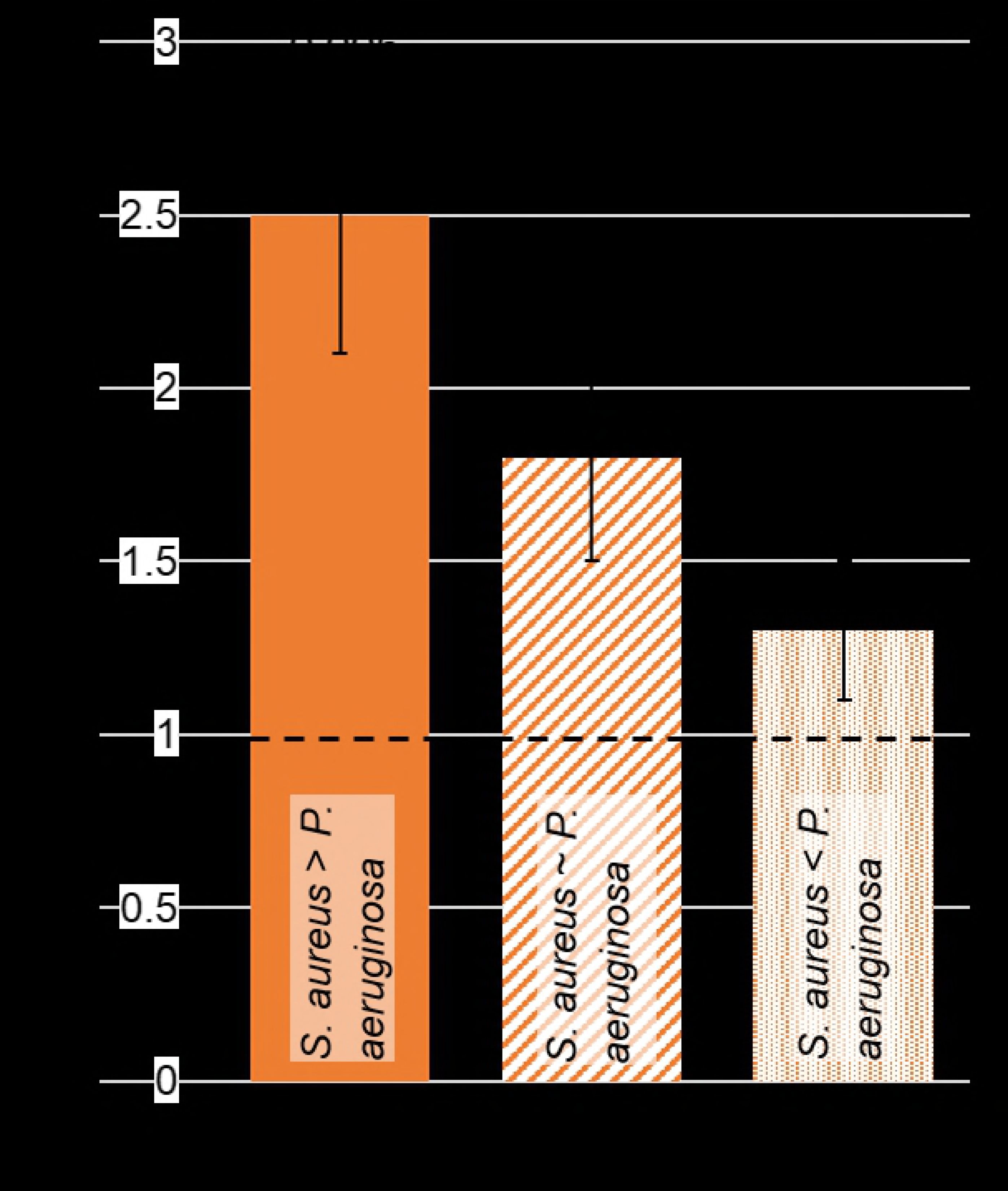
The growth advantage of *P. aeruginosa* aggregates increases when *S. aureus* single cells locally outnumber *P. aeruginosa* single cells. The relative fitness for aggregates increases as the initial local ratio of *S. aureus* to *P. aeruginosa* increases. For each condition, p-values were calculated for the null hypothesis that relative fitness *f* = 1, and are indicated on each bar.

If *S. aureus* were producing a factor harmful to *P. aeruginosa*, from which aggregates are better-protected than single cells, we would expect *P. aeruginosa* growth in co-culture to be slower than growth in monoculture. On the contrary, we find that *P. aeruginosa* in co-culture grows as fast or faster than *P. aeruginosa* in monoculture, for all cases examined (Fig 2).

Thus, our results indicate that *P. aeruginosa* aggregates are able to better obtain some benefit from *S. aureus* than are *P. aeruginosa* single cells, and that the benefit obtained depends on the **<u>local</u>** concentration of *S. aureus*. Because the intrinsic disadvantage of aggregates that we found in our earlier work, namely that cells in the aggregate interiors have very limited access to growth substrate and therefore grow very slowly [22], should still be present in this system, we infer that the benefit obtained from *S. aureus* by aggregates is sufficient to overcome this advantage.

### The extracellular polysaccharide Psl is linked to the growth advantage of *P. aeruginosa* aggregates

Cells in aggregates are bound together by a matrix dominated by extracellular polysaccharides (EPS). Thus, bacteria in aggregates experience a higher local concentration of EPS than do their single-cell counterparts. *In vitro*, the PAO1 lab strain of *P. aeruginosa* makes two types of EPS, Psl and Pel, but Psl is the most dominant [24, 28]. In microscopy observations of a Psl over-expressing strain (Δ*wspF* Δ*pel*) and a Pel over-expressing strain (Δ*wspF* Δ*psl*), we find that shaken liquid cultures of Psl overexpressors than do more and larger aggregates than do wild-type (WT) cultures, whereas shaken liquid cultures of Pel over-expressors have fewer and smaller aggregates than do WT cultures. Therefore, we conclude that Psl is the most important EPS factor for causing *P. aeruginosa* to aggregate.

To investigate the role that Psl may play in the relative fitness of *P. aeruginosa* aggregates and single cells, we performed flow cell coculture experiments with a strain, PAO1 Δ*pel*, which does not produce the EPS Pel. Because there is overlap in the substrates and cellular machinery used for Pel and Psl production [29], we expect that Δ*pel* should make more Psl than the wild-type (WT); this is supported by our earlier study (Fig 6 in [28]). For Δ*pel* aggregates compared with *Apel* single cells, we found a relative fitness of 0.6 ± 0.15 for medium initial density (OD~0.01). Thus, the relative fitness found for WT aggregates vanishes and Δ*pel* aggregates are *less* fit than Δ*pel* single cells. This is true even when the data are analyzed as a function of the local ratios of *P. aeruginosa* and *S. aureus*.

Such a change in relative fitness could arise from an increase in the fitness of the single cells or from a decrease in the fitness of the aggregates. To assess which is occurring here, we measured the absolute fitness (^*N*^/*N*_0_) of single cells and aggregates, for both WT and Δ*pel*. We find that ^*N*^/*N*_0_ is greater for Δ*pel* than for WT for both aggregates and single cells (Fig 5); thus, both aggregates and single cells have increased fitness with increased Psl production. However, the aggregates are 56% fitter with increased Psl while the single cells are 278% fitter with increased Psl. This indicates that the growth advantage of aggregates does not arise from their high density of bacteria, but rather their high Psl content. When single cells express high levels of Psl, they accrue more of the benefits associated with Psl without the disadvantages of being in an aggregate, namely limited access to growth resources in the aggregate interior.

**Fig 5.**
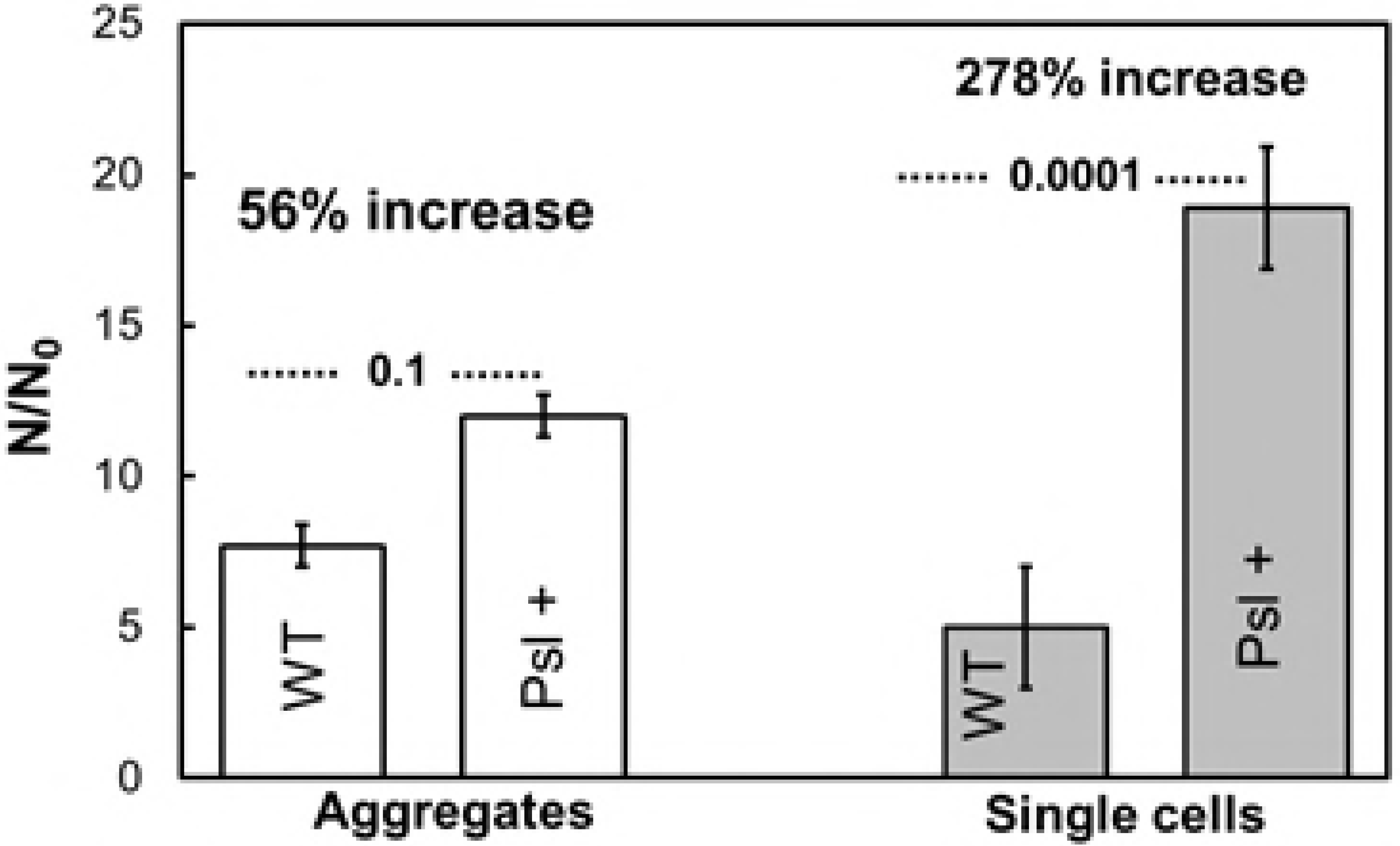
Increased production of Psl (Δ*pel* over-expresses Psl, labeled in figure as Psl+) increases the fitness of both aggregates and single cells in co-culture with *S. aureus*. However, the increase in fitness is greater for single cells than for aggregates - aggregate fitness increases by 56% better, whereas single cell fitnesses increases by 278%. p-values comparing each pair of bars (indicated by dotted lines) are calculated for the null hypothesis that the relative fitness *f* for the paired data are the same.

Thus, the ability to benefit from *S. aureus* that are nearby in space is also associated with a spatially-localized property, the amount of Psl near cells.

### The relative fitness of aggregates in co-culture is linked to low iron content in the environment

Iron is an essential micronutrient for growth of *P. aeruginosa* [30, 31]. All the flow cell experiments presented above were done in M9+glucose, without added iron. Thus, any iron present is an impurity carried over from the rich overnight culture media in which both bacterial species grew. Visaggio *et al.* recently found that multicellular aggregates of *P. aeruginosa* promote pyoverdine-dependent iron uptake into bacterial cells [32]. Yu *et al*. recently found that Psl, which is the primary extracellular polysaccharide promoting aggregation, can bind to and store iron [21]. Thus, there is a link between iron acquisition and both the formation of aggregates and the production of Psl. Recent work has shown that Psl remains closely associated with the cells that produce it [33]. Therefore, we hypothesize that Psl-bound iron may stay with the Psl producer and be more easily acquired and used by that bacterial cell or aggregate.

To determine the effect of sample-wide iron concentration on the relative fitness of *P. aeruginosa* aggregates in co-culture, we performed a series of experiments in which we added iron, in the form of FeSO_4_, to the experimental media in three different concentrations: 0.05, 5 and 100 μ-molar. In each experiment we used a medium initial density of cells (OD~0.01). Compared with no added iron, we find that the relative fitness of aggregates increases when a moderate (5 μ-molar) concentration of iron is added, and increases even more when a low (0.05 μ,-molar) concentration of iron is added; at high concentrations of added iron (100 μ,-molar), aggregates and single cells are equally fit (Fig 6). This suggests that when iron is a scarce resource, aggregates are better able to acquire iron and grow than are single cells, but when iron is abundant this advantage is no longer important.

**Fig 6.**
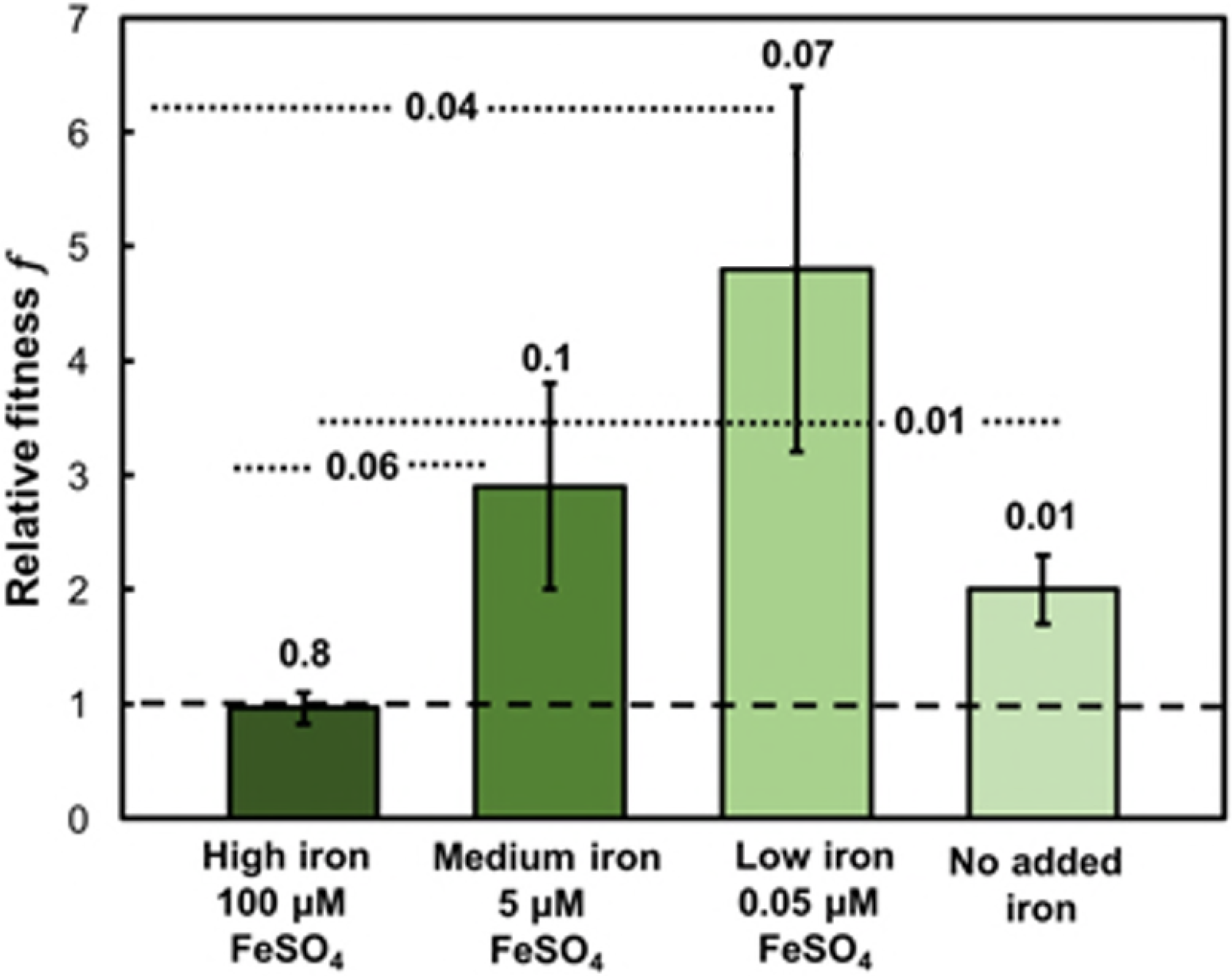
In co-culture with *S. aureus*, the growth advantage of *P. aeruginosa* aggregates over *P. aeruginosa* single cells is modulated by iron content in the growth medium. *p*-values for each single bar are calculated for the null hypothesis that the relative fitness *f* = 1; this is the value indicated by the dashed line and corresponds to no fitness difference between aggregates and single cells. *p*-values comparing each pair of bars (indicated by dotted lines) are calculated for the null hypothesis that the relative fitness *f* for the paired data are the same. These indicate that the differences in relative fitness seen when comparing high-iron conditions with the other conditions tested are statistically significant or, in the case of high *versus* medium added iron, nearly so.

### Reducing iron availability with iron chelator complex increases the relative fitness of aggregates

As another approach to evaluating the importance of iron availability to the relative fitness of aggregates in co-culture, we added an iron chelator, to M9+glucose without explicitly-added iron. EDDHA primarily chelates the ferric form of iron, though it can also chelate small amounts of ferrous iron. By chelation, EDDHA effectively reduces the concentration of iron available to bacteria. Experiments were done at high and medium initial sample-wide cell density (OD~0.1 and ~0.01) with equal amounts of *P. aeruginosa* and *S. aureus* (1:1 as measured by OD). We used two different concentrations of EDDHA - low (32 μ,-molar) and high (320 μ,-molar). For both initial cell densities and both concentrations of chelator, we find that aggregates are fitter than single cells (Fig 7). At low chelator concentration, the relative fitness of aggregates is comparable to that with no chelator present (Fig 2). At high chelator concentration, aggregate advantage over single cells increased by ~90% for high inoculation density, and by ~30% for medium inoculation density (Fig 7).

**Fig 7.**
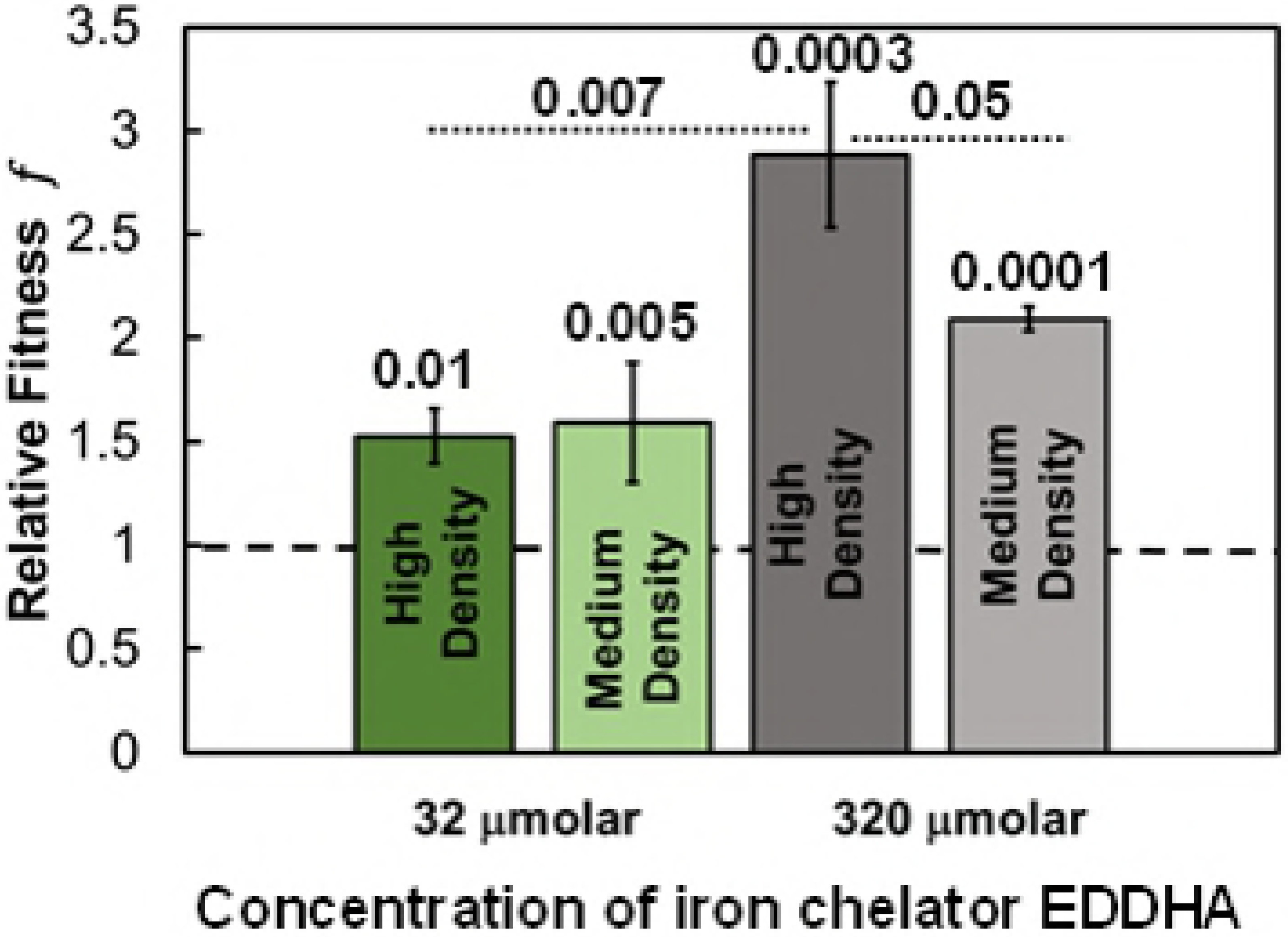
When grown in the presence of high concentrations of iron chelator, the fitness advantage of aggregates increases. P-values shown above single bars were computed using the null hypothesis that the relative fitness *f*=1, meaning no fitness difference between aggregates and single cells. p-values comparing relative fitness under two different conditions are shown above a dotted line indicating the two values being compared, and were computed using the null hypothesis that the relative fitness is the same under both conditions.

Increasing iron chelator is equivalent to decreasing the available iron. Thus, these findings are consistent with aggregates being better able to acquire and use iron for growth when iron is a scarce resource, as we also inferred from the data shown in Fig 6 in the previous subsection. To probe this more deeply, we examine the absolute fitness ^N^/N_0_ changes with iron availability. At high chelator concentration, average ^N^/N_0_ forsingle cells was ~1.1 for high inoculation density and ~2.2 for medium inoculation density; for aggregates, average ^N^/N_0_ was ~3.2 for high inoculation density and ~4.7 for medium inoculation density. Thus, both aggregates and single cells grow more slowly under conditions of high chelator concentration and therefore low iron availability. However, comparing growth under high and low chelator concentration shows that the <u>absolute</u> fitness of single cells is reduced by ~65% when chelator concentration is reduced and the absolute growth of aggregates is reduced by only ~45%. This is the origin of the increase in the <u>relative</u> fitness of aggregates with higher chelator concentration (and therefore lower iron availability). That the <u>absolute</u> fitness of both aggregates and single cells increases with decreasing inoculation density is consistent with our earlier work showing that higher competition for growth resources reduces the rate of growth for all bacteria [22].

### *P. aeruginosa* aggregates do not impair *S. aureus* growth more than do *P. aeruginosa* single cells

Others have shown that aggregation of *P. aeruginosa* leads to increased production of virulence factors by the multicellular aggregate and to quorum sensing that is linked to virulence [25, 32]. These virulence factors are diffusible and allow *P. aeruginosa* to lyse *S. aureus* for use as an iron source [31, 34]. Therefore, it is plausible that the relative fitness advantage of aggregates could arise, at least in part, from aggregates producing more virulence factors than single cells.

To evaluate whether *P. aeruginosa* aggregates are associated with increased lysis of *S. aureus* in the local vicinity, we examined how fitness of *S. aureus* depends on its proximity to a multicellular aggregate of *P. aeruginosa*. This analysis is conceptually similar to the analysis done to determine the relative fitness of *P. aeruginosa* aggregates with respect to *P. aeruginosa* single cells. We calculated the fractional increase in *S. aureus* biomass *N*_Staph_/*N*_Staph_0_ for *S. aureus* near the *P. aeruginosa* aggregates 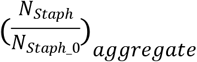 and ~500 μm away 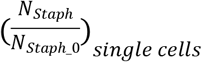. The relative fitness *f*_*staph*_ was 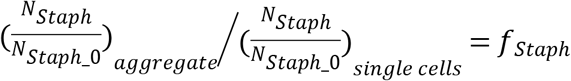.

Surprisingly, we found that *S. aureus* fitness is unchanged by proximity to an aggregate of *P. aeruginosa* (Fig 8). This was true for all initial cell densities, and all initial ratios of the two species, except at moderate inoculation density where the *S. aureus* fitness is actually <u>higher</u> next to a *P. aeruginosa* aggregate. These results imply that multicellular aggregates do not have an advantage in lysing nearby *S. aureus* and thereby releasing iron.

**Fig 8.**
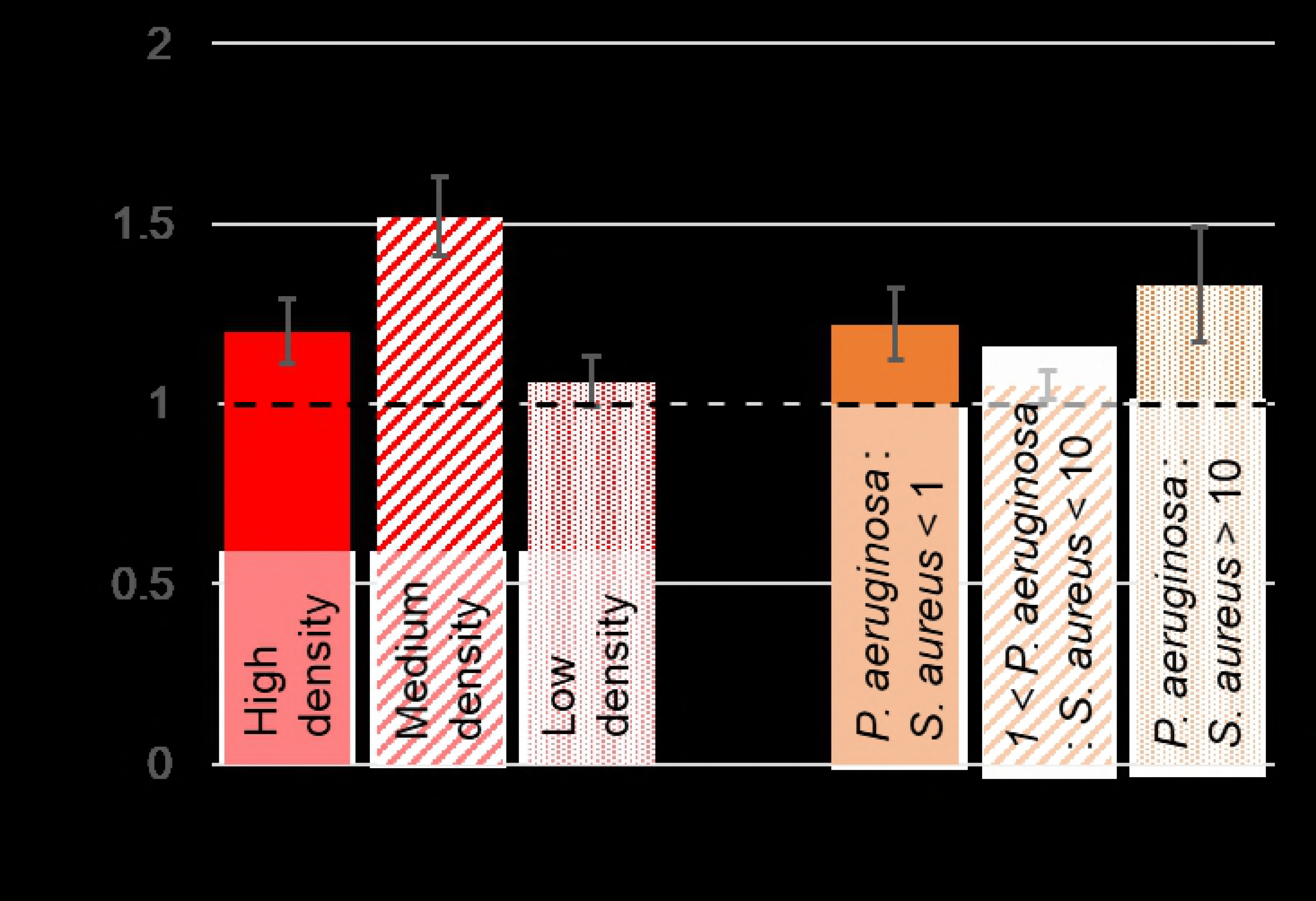
The fitness of *S. aureus* cells near a *P. aeruginosa* aggregate, compared with *S. aureus* cells far from an aggregate. (but near *P. aeruginosa* single cells) is not significantly impacted by the inoculation density or by the local ratio of the two species. P-values were calculated for the null hypothesis that relative fitness of *S. aureus* is 1, and are indicated above each bar.

Since killing of *S. aureus* by *P. aeruginosa* does not appear linked to the relative fitness of *P. aeruginosa* aggregates, we also investigated whether the benefit provided by *S. aureus* requires living activity on the part of *S. aureus*. For this, we killed S. aureus using heat and then inoculated, imaged, and measured the relative fitness of P. aeruginosa aggregates as before. Under these conditions, the relative fitness of P. aeruginosa is reduced and is no longer statistically significant (Fig 9) This suggests that active production and release of a factor by *S. aureus* is important for conferring a fitness advantage to *P. aeruginosa* aggregates [35–37].

**Figure 9.**
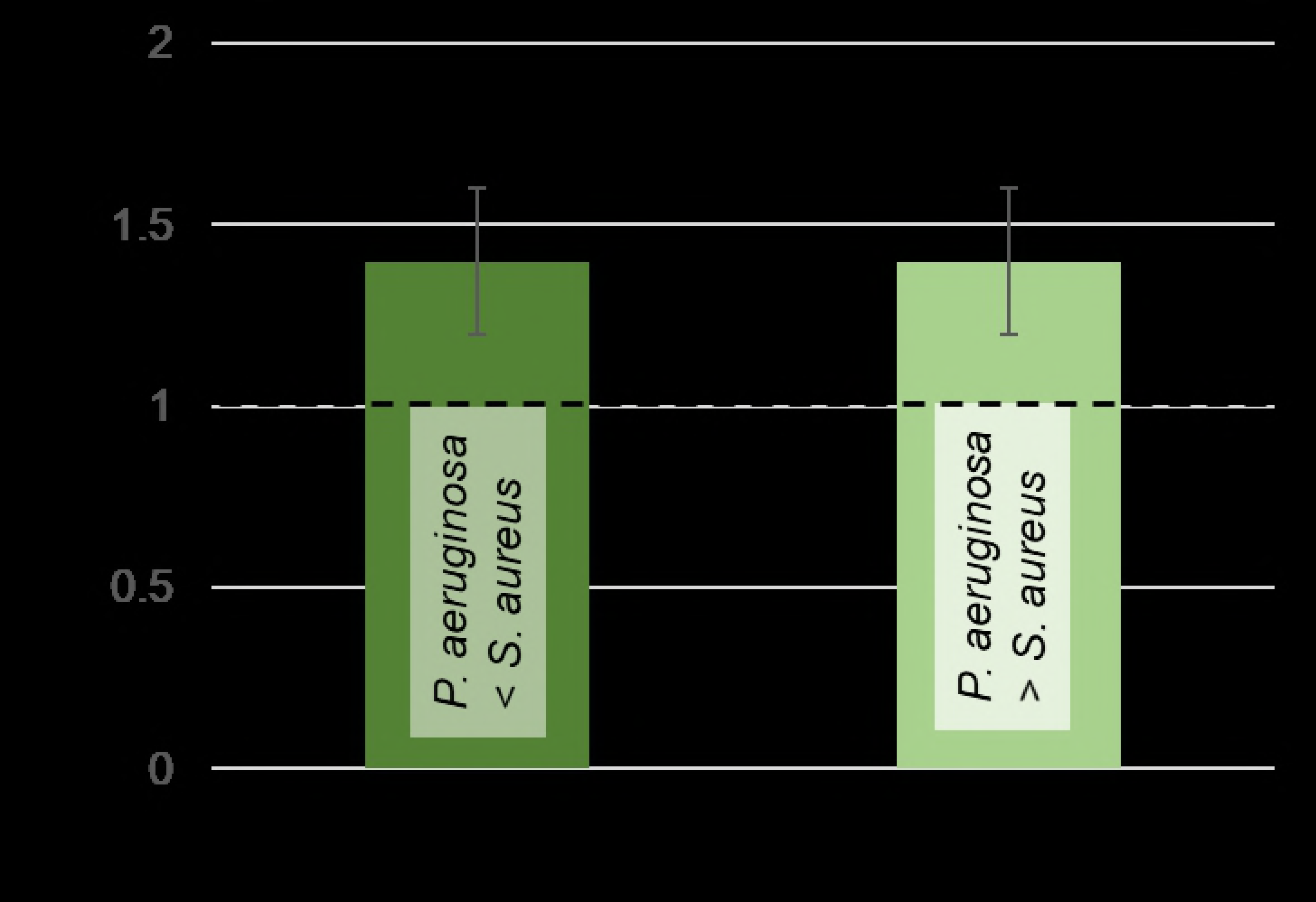
The fitness advantage of *P. aeruginosa* aggregates is reduced when *S. aureus* has been heat-killed before inoculation. P-values are calculated using the null hypothesis that aggregates grow at the same rate as *P. aeruginosa* single cells. When all the *S. aureus* present have previously been heat-killed, the growth of aggregates is no longer statistically-significantly different from that of single cells, and the impact of local ratio of *P. aeruginosa* to *S. aureus* is no longer seen.

## Discussion

### Hypothesis: The *P. aeruginosa* EPS Psl binds siderophores produced by *S. aureus*

Thus, we conclude that *P. aeruginosa* aggregates are better able than *P. aeruginosa* single cells to utilize a benefit that matters when the availability of iron is low, that originates from *S. aureus*, and that increases with the local density of *S. aureus*. However, the disproportionate benefit to aggregates is not due to aggregates being more adept at killing nearby *S. aureus*, and likely requires the production of an active factor by S.*aureus. S. aureus* is known to produce siderophores that bind iron [35–37]. Therefore, we hypothesize that *P. aeruginosa* aggregates are better able to acquire iron from *S. aureus* siderophores than are *P. aeruginosa* single cells, and that this may be linked to the ability of the EPS Psl to bind iron. This hypothesis will be the subject of future investigation, with further considerations described below.

### Types of iron available

The growth advantage of *P. aeruginosa* aggregates over *P. aeruginosa* single cells is linked to low availability of iron in the environment. FeSO_4_ adds the ferrous form of iron (iron II), which is soluble in water. However, at moderate and high levels of added iron, there is obvious precipitate in the experimental media. Iron II is spontaneously oxidized to iron III (ferric form) by reacting with molecular oxygen (present in aqueous media). Ferric iron is insoluble in water and likely causes the precipitates we observe. Therefore, the growing biofilm probably has access to a mixture of soluble iron II (ferrous form) and small quantities of insoluble iron III (ferric form). *P. aeruginosa* has different strategies for acquiring different forms of iron: it uses the Feo system to uptake soluble ferrous iron [38], and it uses siderophores to uptake ferric iron [39]. Both systems could be in use in these experiments. Future work could distinguish Psl binding directly to iron from the benefits conferred by *S. aureus* siderophores by using *S. aureus* siderophore mutants or staphyloferrin.

### Relative fitness likely results from an interplay between aggregate structure and Psl content

The finding that increased Psl production benefits single cells more than aggregates (Fig 5) likely arises from the intrinsic structural disadvantage of aggregates, namely that cells in the interior grow slowly because they have limited access to growth substrate [22]. Single cells do not have this disadvantage.

### Implications for the evolutionary development of biofilms

Iron scarcity and the presence of multiple species are widespread characteristics of the natural environments in which *P. aeruginosa* evolved. The work we present here suggests that the iron-binding properties of Psl may have provided a selective advantage that also led to greater tendencies to aggregation. Such aggregative tendencies could also have promoted the development of additional group benefits associated with *P. aeruginosa* aggregates and biofilms [25]. Because we find that the iron-associated growth benefit for aggregates does not depend on the density of *P. aeruginosa* competitors for growth resources, there is no reason to believe that spatial structure is a major driver for this interaction. This is strikingly unlike our earlier finding for *P. aeruginosa* monoculture, in which competition and the spatial structure of both aggregates and the environment are essential to the growth advantage for aggregates [22]. Releasing the requirement for spatial structure greatly broadens the range of natural scenarios in which the aggregate growth advantage could provide a selective advantage on which evolution might act.

### Implications for biofilm disease

In addition to being a more realistic representation of natural environments, polymicrobial systems account for most pathogenic infections [40–44]. *P. aeruginosa* and *Staphylococcus aureus* are widespread co-pathogens in wounds, catheters, implants, and in the cystic fibrosis lung [45–57]. Interspecies interactions, including those between *P. aeruginosa* and *S. aureus*, can strongly impact clinical outcomes [45, 48, 58–67]. Therefore, better understanding of the ecology of *P. aeruginosa* + *S. aureus* systems is likely to lead to better treatments of polymicrobial infections.

For instance, our work suggests that high Psl production could contribute to fitness by promoting iron acquisition. This might contribute to the evolutionary trend toward higher Psl production that is recently being recognized for chronic infections in cystic fibrosis [68–71]. We have recently suggested that Psl may confer a mechanical fitness benefit on biofilm infections, by increasing their elasticity, yield stress, and toughness, and perhaps thereby hindering breakup by phagocytosing neutrophils [28]. Better iron acquisition would be a chemical and metabolic advantage of Psl production that complements and is orthogonal to its effects in increasing mechanical robustness.

## Conclusion

Our results show that aggregates of *P. aeruginosa* can have a growth advantage over single cells in environments with limited iron. This growth advantage is linked to Psl, the dominant cohesive EPS component of PA01 aggregates; when single cells overexpress Psl, their growth becomes comparable to that of aggregates. Based on the recent work of others [21], we suggest that this is likely because Psl recruits iron so that cells embedded in high-Psl environments experience a resulting enrichment of environmental iron as well.

## Acknowledgements

This work was funded by: grants from the Human Frontier Science Program (HFSP-RGY0081/2012 - GORDON), the Cystic Fibrosis Foundation (Gordon 201602808-001), the National Science Foundation (727544, BMMB, CMMI), and the National Institutes of Health (1R01AI121500-01A1, NIAID), all to Vernita Gordon; a gift from ExxonMobil; startup funds from The University of Texas at Austin. Professor Shelley Payne (The University of Texas at Austin) gave us deferrated EDDHA and read and commented on the manuscript before submission. Professor Marvin Whiteley (The University of Texas at Austin) gave us the plasmid pJY209.

## Supporting information

**S1 Fig. *S. aureus* monocultures grow very poorly compared with *P. aeruginosa* monocultures under the same conditions.** This indicates that *S. aureus* does not strongly compete with *P. aeruginosa* for growth resources. (A) In shaken liquid culture over 40 hours of growth after inoculation, *P. aeruginosa* cultures attain optical densities ~4-5 times higher than corresponding *S. aureus* cultures. Optical density is roughly proportional to the density of bacteria. (B) In flow cells in monoculture, the average number density of *S. aureus* slowly declines. Error bars are standard error of the mean.

